# GRABS: a content-addressed antibody-antigen binding-affinity dataset with predicted structures and a leak-free benchmark

**DOI:** 10.64898/2026.11.17.767263

**Authors:** Luciano Martins, Priscila Pini Zenatti, João Meidanis

## Abstract

Machine-learning prediction of antibody-antigen binding affinity depends as much on the training data as on the model, yet public affinity measurements are scattered across databases with incompatible formats, mix genuine dissociation constants with censored or predicted values, and concentrate most of their label mass on a few heavily mutated targets. We present **GRABS** (*Geometric Resource for Antibody Binding Studies*), a curated, **content-addressed**, structure-ready dataset that consolidates nine public sources, of which the seven with redistributable licenses yield **4**,**380 canonical antibody-antigen records** (median-collapsed from **4**,**761 raw measurements**), each with an **experimentally measured dissociation constant (***K*_*d*_), paired heavy and light variable regions (VH, VL), and an antigen sequence, covering **2**,**015 distinct antigen sequences**. Every structure, graph, and feature is addressed by a content key, `md5(VH+VL+antigen)[:10]`, and verified non-circularly: each graph’s node features are decoded back to sequence and re-hashed, matching the content key for **4**,**282/4**,**282 (100%)** structure-backed records. This integrity is load-bearing: an alignment ablation shows that a previously-reported strong cross-target result (*R*^2^ *≈* 0.415) was a misalignment-mediated source/batch leak that content-key regeneration dissolves. On the resulting leak-free, antigen-cluster-grouped benchmark (4,020 records), cross-target ranking of variants of a held-out antigen collapses to the noise floor (within-held-out-antigen Spearman *ρ ≈* 0 for every model family: sequence, structure, and folding-confidence), with structure giving no advantage, a rigorously leakage-controlled quantification of the limited reach of current methods. GRABS ships model-ready artifacts (Boltz-2 predicted complexes, per-residue ESM-2 embeddings, geometric graphs, feature vectors) with antigen-stratified splits, and is intended for leakage-free benchmarking and within-target affinity ranking (measure-then-rank), not zero-shot prediction for antigens absent from training.

## 1 Background & Summary

Monoclonal antibodies are among the fastest-growing classes of biopharmaceuticals, supported by an expanding therapeutic pipeline and continued advances in antibody discovery and engineering [1, 2]. The property that engineering campaigns most often predict and optimize is binding affinity, quantified by the dissociation constant *K*_*d*_ (related to the standard binding free energy by Δ*G*^*°*^ = *RT* ln *K*_*d*_) [3, 4]. Data-driven models for affinity, from gradient-boosted trees to geometric graph neural networks [5, 6], depend heavily on the quality and composition of the training corpus [7]. Yet public affinity data remain fragmented across databases with incompatible schemas, units, and provenance, and often mix quantitative experimental measurements with qualitative, binary, or computationally predicted labels.

Three problems follow. The first is **evaluation leakage**: when splits are not made by antigen, models memorize target-specific signal and report optimistic numbers that say little about generalization to new targets, a failure mode documented for small-molecule ligands [8] and equally acute here [9]. The second is **target concentration**: a few antigens subjected to deep mutational scanning dominate the label mass, biasing both training and naive evaluation. The third, and the one that motivates our content-addressed design, is **silent feature-label misalignment**: when derived features (embeddings, graphs) are linked to records by positional index, a re-sort or re-index can pair a feature with the wrong label, and the resulting artifact can masquerade as predictive signal.

GRABS addresses these directly. It unifies nine sources into a single schema with normalized *K*_*d*_ and full provenance; applies an explicit, auditable curation pipeline that removes contaminated and censored records and rebalances over-represented targets; and ships model-ready artifacts with antigen-stratified, leakage-free splits. Distinctively, it is **content-addressed**: every record and its artifacts are keyed by a hash of the record’s own sequences, so a feature can never silently desynchronize from its label, and the integrity is verified by decoding each graph back to sequence and re-hashing. We show that this guardrail is not cosmetic: it dissolves a strong but spurious cross-target result present in an earlier positionally-indexed build. We then report the honest downstream picture, including the difficulty of predicting affinity for unseen antigens, and position GRABS for researchers who need a standardized, integrity-verified basis for benchmarking affinity predictors and for measure-then-rank affinity maturation.

## 2 Methods

### 2.1 Source aggregation

GRABS consolidates nine public antibody-antigen affinity datasets (eight databases, with SAbDab contributing two subsets). Each source was parsed into a common schema comprising VH, VL, antigen sequence, a numeric *K*_*d*_ in molar units, and a provenance tag. Sources contributing qualitative, binary, or purely predicted affinities were excluded at ingestion; only records traceable to an experimental measurement (SPR, BLI, ITC, ELISA, or curated literature) were retained. Seven sources permit redistribution and are shipped here; two (AgAb/ASD [10], non-commercial; AbCoV [11], no explicit reuse license) are provided as source pointers rather than redistributed data (Section 3). Table 1 lists the composition at both the raw-measurement level (`measurements.csv`, pre-deduplication) and the canonical level (`grabs.csv`, after median-log deduplication).

**Table 1:** Composition of the redistributed GRABS release by source, at the raw-measurement and canonical levels, spanning 2,015 distinct antigen sequences. A source’s canonical count can differ from its raw contribution because of median-log deduplication of (VH, VL, antigen) triples, the quality-controlled ANDD re-extraction, and re-attribution of re-hosted measurements. Two further sources whose licenses do not permit redistribution (AgAb/ASD, 316; Ab-CoV, 11; 327 records total) are provided as source pointers, not raw data (Section 3).

| **Source** | **Raw meas.** | **Canonical** | ***K*<sub>*d*</sub> measurement** | **Reference** |
|---|---|---|---|---|
| ANDD | 2,311 | 2,023 | Experimental (mixed) | [12] |
| AbRank | 681 | 681 | Experimental (SKEMPI-v2 scan) | [9] |
| SAbDab-ΔG | 566 | 482 | SPR/BLI/ITC/ELISA | [13] |
| AbSci (anti-HER2) | 465 | 465 | SPR | [14] |
| FLAb | 374 | 374 | SPR/BLI | [15] |
| SAbDab (Zenodo subset) | 342 | 333 | SPR/BLI/ITC/ELISA | [16, 13] |
| CoV-AbDab | 22 | 22 | SPR/BLI | [17] |
| **Total (redistributed)** | **4,761** | **4,380** | **100% experimental** |  |

### 2.2 Curation pipeline

On top of aggregation, GRABS applies a documented, reproducible curation pipeline (Figure 1). *K*_*d*_ values are normalized to molar units and stored on a log_10_ scale, with values outside a biologically plausible window (10^*−*15^ to 10^−2^ M, informed by the physical limits of antibody affinity [18]) discarded. VH and VL are validated and IMGT-renumbered with ANARCI [19]; records with non-standard amino acids or failed numbering are removed. Records whose declared antigen is in fact an antibody variable domain (a recurrent labeling error in aggregated sources) are removed by decontamination, and records with placeholder or threshold *K*_*d*_ values (exact repeated detection-limit constants incompatible with genuine measurements) are removed by de-placeholding. The deep-mutational-scanning source (AgAb/ASD, dominated by a single influenza-hemagglutinin campaign) is down-sampled by *K*_*d*_-stratified sampling so that no single target dominates, while curated SPR/ranking campaigns (for example the AbSci anti-HER2 set and AbRank) are retained uncapped. Repeated (VH, VL, antigen) triples are collapsed into a single record whose *K*_*d*_ is the median of the constituent log_10_ *K*_*d*_ values, and a SAbDab-ΔG structural pack is added for antigen diversity.

**Figure 1.**
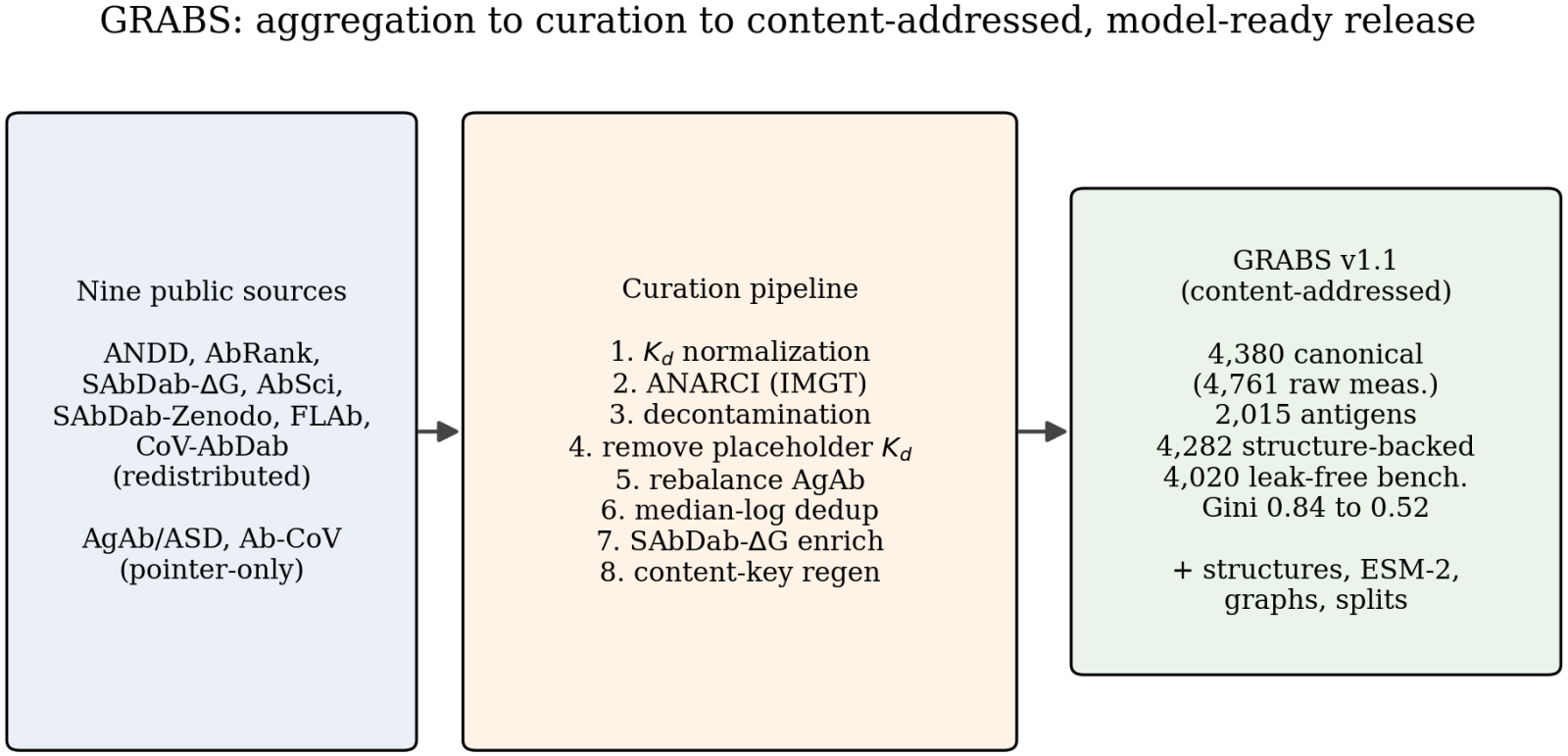
Construction of GRABS. Nine public sources (seven redistributed, two provided as pointers) pass through the curation pipeline (*K*_*d*_ normalization, ANARCI IMGT numbering, decontamination, placeholder removal, *K*_*d*_-stratified target rebalancing, median-log deduplication, structural enrichment, and content-key regeneration) to the content-addressed v1.1 release: 4,380 canonical records (from 4,761 raw measurements) over 2,015 antigens, of which 4,282 are structure-backed and 4,020 form the leak-free benchmark, with the per-antigen Gini falling from 0.84 to 0.52.

The net effect is a well-distributed corpus: the per-antigen entry distribution has a Gini coefficient of **0.52** (down from 0.84 in the raw pool), and the most frequent antigen, HER2, accounts for **16.6%** of records (729 of 4,380), down from a single influenza-hemagglutinin strain at 52% of the raw pool. All retained *K*_*d*_ values are experimental; there are no placeholder decade constants and no duplicate (VH, VL, antigen) triples. Of the 4,380 canonical records, 3,699 are identity-level measured pairs and 681 are flagged `label type = skempi mutational scan` (within-complex mutational-scan data from AbRank’s quantitative subset, a distinct data type).

Building the dataset also exposed a limit of the field: even after pooling every accessible public source, no corpus of quantitative *K*_*d*_ is simultaneously large, clean, and broadly target-diverse. Sources with many antigens tend to carry weak, binary, or predicted labels, while sources with clean *K*_*d*_ cover few targets. This label-diversity ceiling is one reason cross-target prediction remains hard (Section 4.3), and GRABS documents it openly.

### 2.3 Content addressing and integrity verification

Every structure, graph, and feature vector is addressed by a **content key**

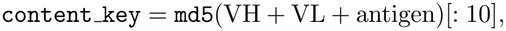

and artifacts are named by this key (`structures/<content key>.pdb`, `graphs/<content key>.pt`), with `features/global features.npz` carrying a parallel `content keys` array. Because the key is a hash of the record’s own content, a feature can never be silently paired with the wrong label by a re-sort or re-index.

This is verified *non-circularly*. Each graph’s per-residue node channels (amino-acid identity and chain) are decoded back to sequence and re-hashed; the result matches the content key for **4**,**282/4**,**282 (100%)** structure-backed records. The ESM-2 node and global features were additionally confirmed faithful by re-embedding the sequences independently (cosine similarity 1.000). Integrity is therefore a property of the shipped files, not a claim.

### 2.4 Derived structural and learned features

Each structure-backed record is augmented with derived artifacts. Three-dimensional antibody-antigen complexes are predicted with Boltz-2 [20] (training cutoff 2023-06-01), retaining per-complex confidence metrics (iPTM, PTM, pLDDT). Per-residue representations (1,280-dimensional) are extracted from the ESM-2 protein language model [21] (`esm2 t33 650M UR50D`, layer 33). From the predicted complexes we build geometric graphs with GVP-style features [5], where nodes are residues (carrying the per-residue ESM-2 embedding, region and chain identity, and solvent accessibility) and edges encode sequential and spatial relationships, together with antibody and antigen masks. A 2,616-dimensional per-complex feature vector (ESM-2 means plus physicochemical descriptors) is provided in `features/global features.npz`. Of the 4,380 canonical records, **4**,**282** are structure-backed (a Boltz-2 complex, a graph, and a feature vector exist, flagged `has structure = True`); the remaining 98 are metadata-only.

### 2.5 Antigen clustering and leak-free splits

Because generalization to *new targets*, not merely new variants of known targets, determines practical value [8, 9], GRABS ships antigen-stratified splits. Antigen sequences are clustered with MMseqs2 [22] (minimum sequence identity 0.8, coverage 0.8); the 4,282 structure-backed records fall into **582 clusters**. A five-fold cross-validation grouped by cluster defines the **4**,**020-record leak-free benchmark** (`in benchmark = True`; five balanced folds of 804 records), which spans **1**,**752 antigens in 549 clusters**. Grouping by cluster ensures that near-duplicate antigens (for example SARS-CoV-2 spike and its RBD) are always held out together, so no antigen-specific signal leaks between train and test.

## 3 Data Records

GRABS is deposited in Zenodo (Data Citation 1) as a content-addressed bundle. The canonical table `grabs.csv` holds the 4,380 records, each row carrying its `content key`, VH, VL, antigen, *K*_*d*_ (molar and log_10_), IMGT CDR annotations, `label type`, `antigen cluster`, `cv fold`, and the boolean flags `has structure` and `in benchmark`. Artifacts are joined to rows by `content key`, never by row order. Records from the two non-redistributable sources are listed as pointers in `restricted records.csv` (a per-source summary covering 327 records: AgAb/ASD 316, Ab-CoV 11); `FETCH RESTRICTED.md` documents how to obtain them and rebuild their artifacts with the released pipeline. Table 2 summarizes the release.

**Table 2:**
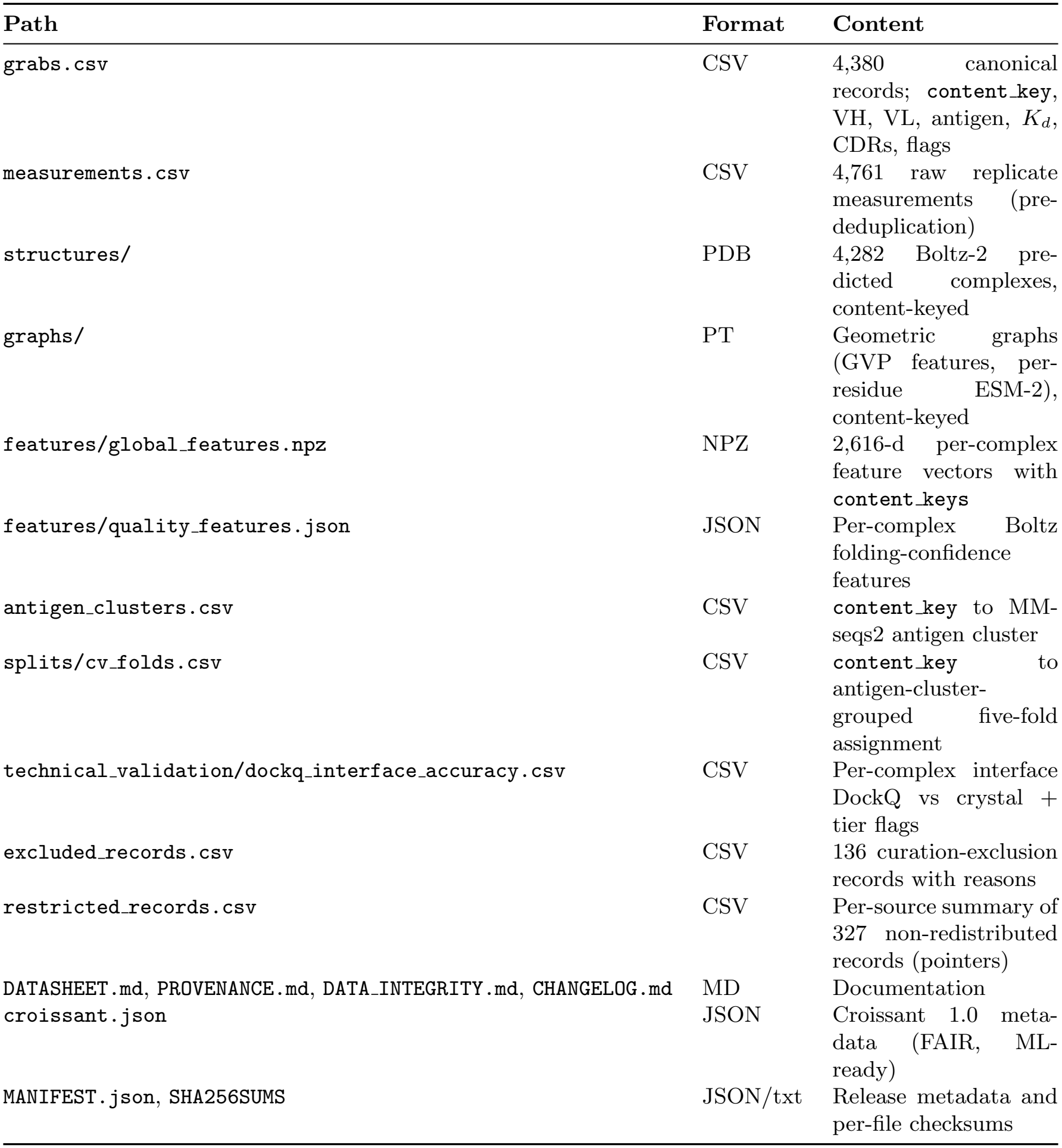
Structure of the GRABS v1.1 release. Every per-record artifact is named by content key; has structure flags the 4,282 structure-backed records and in benchmark flags the 4,020 leak-free benchmark records.

| **Path** | **Format** | **Content** |
|---|---|---|
| grabs.csv | CSV | 4,380 canonical records; content_key, VH, VL, antigen, *K<sub>d</sub>*, CDRs, flags |
| measurements.csv | CSV | 4,761 raw replicate measurements (pre-deduplication) |
| structures/ | PDB | 4,282 Boltz-2 pre-dicted complexes, content-keyed |
| graphs/ | PT | Geometric graphs (GVP features, per-residue ESM-2), content-keyed |
| features/global_features.npz | NPZ | 2,616-d per-complex feature vectors with content_keys |
| features/quality_features.json | JSON | Per-complex Boltz folding-confidence features |
| antigen_clusters.csv | CSV | content_key to MM-seqs2 antigen cluster |
| splits/cv_folds.csv | CSV | content_key to antigen-cluster-grouped five-fold assignment |
| technical_validation/dockq_interface_accuracy.csv | CSV | Per-complex interface DockQ vs crystal + tier flags |
| excluded_records.csv | CSV | 136 curation-exclusion records with reasons |
| restricted_records.csv | CSV | Per-source summary of 327 non-redistributed records (pointers) |
| DATASHEET.md, PROVENANCE.md, DATA_INTEGRITY.md, CHANGELOG.md | MD | Documentation |
| croissant.json | JSON | Croissant 1.0 meta-data (FAIR, ML-ready) |
| MANIFEST.json, SHA256SUMS | JSON/txt | Release metadata and per-file checksums |

## 4 Technical Validation

### 4.1 Integrity: a dissolved cross-target leak

The content-addressing is load-bearing, and we validate it adversarially. An earlier, positionally-indexed build of this corpus reported a strong pooled cross-target result (*R*^2^ *≈* 0.415). An alignment ablation, holding the rows, the split, and the labels fixed and toggling *only* the feature store, showed this to be a **misalignment-mediated source/batch leak**: an antigen-length field left unpopulated for five of seven sources had become a source indicator correlated with *K*_*d*_ (correlation 0.59, versus 0.003 for true antigen length), and the antigen-cluster split does not hold out source. Regenerating features in structure-residue order and re-keying by content hash dissolves the inflation entirely (pooled *R*^2^ *≈ −* 0.2, pooled Spearman *ρ ≈* 0.21; within-held-out-antigen ranking *ρ ≈* 0). Content addressing is what makes this class of bug impossible to reintroduce, and the decode-and-rehash check (Section 2.3) makes the guarantee auditable by any user.

### 4.2 Comparison to existing resources

Table 3 positions GRABS against widely used antibody-antigen affinity resources. Mutational databases such as SKEMPI 2.0 [23] and AB-Bind [24] provide changes in binding free energy (ΔΔ*G*) upon point mutation against a fixed reference complex, rather than absolute affinity across many antigens. SAbDab [13] is centered on solved structures with sparse, heterogeneous affinity annotations. Ranking benchmarks such as AbRank [9] supply pairwise order rather than absolute *K*_*d*_. Specialized collections (FLAb [15], CoV-AbDab [17]) are narrow in antigen scope. The closest peer is ANDD [12], a unified antibody-and-nanobody design resource that is itself one of our inputs. GRABS is complementary: it is antibody-focused and affinity-first (censored and qualitative labels removed), it attenuates target concentration, it is content-addressed and integrity-verified, and it ships a model-ready stack (per-residue ESM-2, geometric graphs, antigen-stratified splits) packaged for leakage-free cross-target evaluation. Because several comparators (SAbDab, AbRank, FLAb, CoV-AbDab, ANDD) are also GRABS inputs, the comparison is between resource designs rather than strictly disjoint record sets.

**Table 3:** GRABS relative to existing antibody-antigen affinity resources. The *K*_*d*_ column marks a unified corpus of absolute experimental dissociation constants, as opposed to relative ΔΔ*G*, pairwise ranking, or mixed labels. *Abbreviations:* Ag seq, antigen sequence provided; Struct., experimental complex structure available. Coverage: ✓ full, “partial” incomplete, *×* absent.

| **Resource** | **Scale** | **Label type** | **Ag seq** | **Struct.** | *K<sub>d</sub>* |
|---|---|---|---|---|---|
| SAbDab [13] | ∼thousands structures | structure (+sparse aff.) | ✓ | ✓ | × |
| SKEMPI 2.0 [23] | 7,085 mutations | ΔΔ*G* (mutation) | ✓ | ✓ | × |
| AB-Bind [24] | 1,101 mut. / 32 cplx | ΔΔ*G* (mutation) | partial | ✓ | × |
| AbRank [9] | ranking pairs (derived) | pairwise ranking | ✓ | × | × |
| FLAb [15] | few campaigns | fitness / binding | partial | × | × |
| CoV-AbDab [17] | SARS-CoV-2 antibodies | mixed binding evidence | partial | × | × |
| ANDD [12] | Ab + nanobody design | seq / struct / affinity | ✓ | ✓ | mixed |
| **GRABS** | **4,380 / 2,015 Ag** | **experimental** ***K<sub>d</sub>*** | ✓ | ✓ | ✓ |

### 4.3 Leak-free benchmark: the honest cross-target picture

On the content-key-verified benchmark (antigen-cluster five-fold; 4,020 records, 1,752 antigens, 549 clusters), cross-target affinity prediction is **weak for every model family we tried**. A sequence-only baseline, a structure-based geometric graph neural network, and a folding-confidence baseline all collapse to the noise floor at ranking variants of a held-out antigen: well-powered within-held-out-antigen Spearman *ρ ≈* 0 (offset-removed *R*^2^ *<* 0), with the structure-based model giving **no advantage** over the sequence baseline. A pooled Spearman is higher (*≈* 0.2) only because a model can predict each antigen’s mean *K*_*d*_ offset, not because it ranks variants within a new target. Holding out a whole data source (source leave-one-out) fails outright, because sources are not exchangeable. These numbers are not a defect of GRABS; they are what a leakage-free protocol reveals once the misalignment leak of Section 4.1 is removed, and they are consistent with a companion characterization study on the same release. GRABS makes this regime measurable rather than hidden (Figure 2).

**Figure 2.**
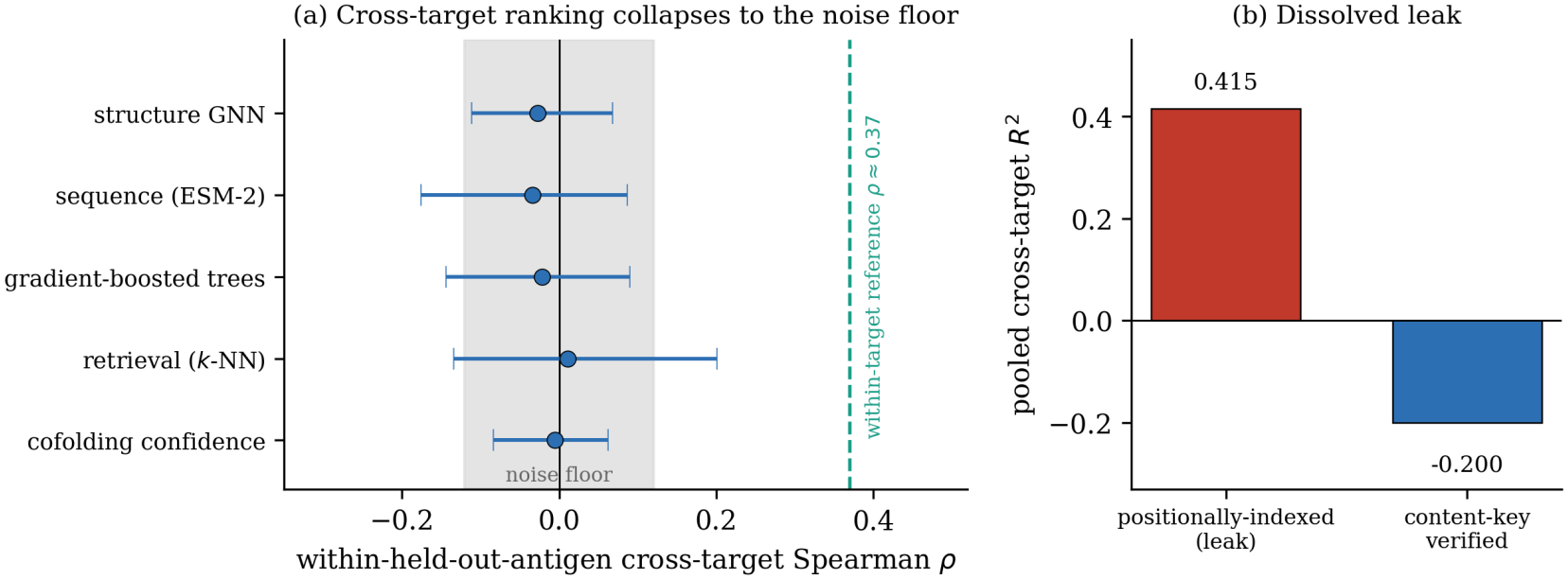
Leak-free benchmark on the v1.1 release. (a) Within-held-out-antigen cross-target Spearman *ρ* for five method families (structure GNN, sequence, gradient-boosted trees, retrieval, and cofolding confidence), all within the noise-floor band at *ρ ≈* 0 with overlapping 95% confidence intervals and far below the within-target measure-then-rank reference (*ρ ≈* 0.37); predicted structure gives no advantage. (b) The dissolved leak: the pooled cross-target *R*^2^ of 0.415 on the earlier positionally-indexed feature store collapses to − 0.20 on the content-key-verified store, confirming the earlier result was a misalignment-mediated artifact.

### 4.4 Predicted-structure interface accuracy

The shipped 3D complexes are Boltz-2 predictions, not experimental structures, and their interface accuracy should be filtered by folding confidence for interface-sensitive work. We quantify this with per-complex interface DockQ against crystal structures where available (`dockq interface accuracy.csv`, 1,726 complexes). The overall median DockQ is **0.766**, but this is contaminated-optimistic: on genuinely novel complexes (released after the Boltz-2 training cutoff, size- and viral-matched) the median collapses to **0.072** (52 complexes). Users relying on the graphs for interface-level analysis should treat structural features as predicted and gate on the shipped confidence metrics.

### 4.5 Antigen diversity and distributional sanity

GRABS is broadly distributed across targets: the per-antigen entry distribution over 2,015 antigen sequences has a Gini coefficient of 0.52, and the most frequent antigen (HER2) accounts for 16.6% of records, a marked reduction from the raw pool where a single influenza-hemagglutinin strain accounted for 52% (Figure 3). All *K*_*d*_ values lie within the plausible [10^−15^, 10^−2^] M window, every record has paired VH/VL and an antigen sequence, and no exact (VH, VL, antigen) triple appears more than once. Residual skew remains: viral antigens (influenza hemagglutinin, SARS-CoV-2) and the anti-HER2 campaign are over-represented relative to the long tail of single-measurement targets, reflecting where quantitative *K*_*d*_ has historically been generated.

**Figure 3.**
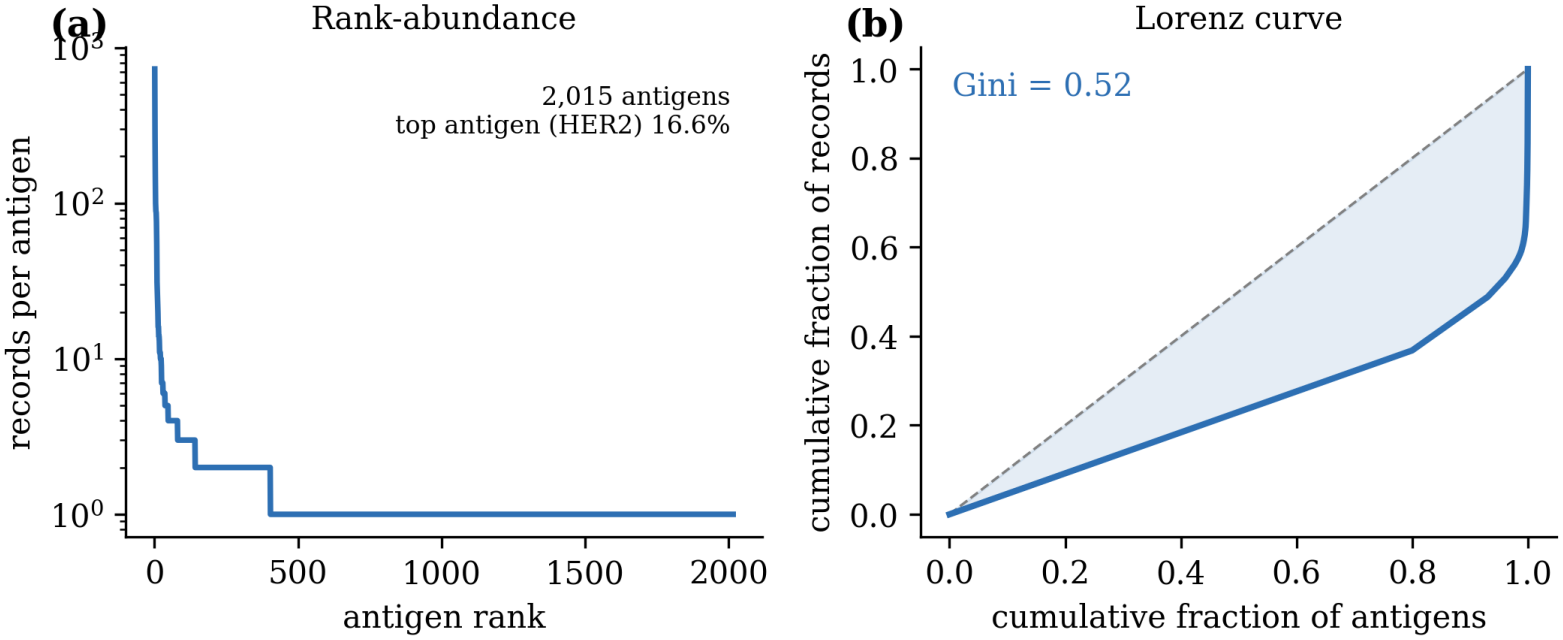
Antigen diversity of GRABS after curation. (a) Rank-abundance (log scale) of perantigen record counts across the 2,015 antigen sequences. (b) Lorenz curve of the same distribution (Gini 0.52); the most frequent antigen, HER2, accounts for 16.6% of records, down from a single influenza-hemagglutinin strain at 52% of the raw pool.

### 4.6 Scope and intended use

GRABS is built to support generalization testing to *new targets*. Under antigen-grouped evaluation, ranking variants of an *unseen* antigen is weak (Section 4.3), whereas within-target ranking, new variants of a target that already has measurements, remains informative. GRABS is therefore intended for within-target affinity ranking (for example measure-then-rank affinity maturation) and for leakage-free benchmarking, and should not be used for zero-shot affinity prediction for antigens absent from training. This within-target versus cross-target boundary is a general feature of antibody-antigen affinity prediction, characterized in a companion study on this release.

## 5 Usage Notes

### Select through the shipped columns

Load `grabs.csv` and join artifacts by `content key`; do not iterate raw files on disk or assume row order. Restrict graph and feature access to rows with `has structure = True` (4,282), and take the benchmark universe as `in benchmark = True` (4,020), never larger.

### Split by antigen, never by record

Use `splits/cv folds.csv` (or `antigen clusters.csv`) so that near-duplicate antigens are always held out together, and verify that no antigen cluster appears in both train and test. Naive random splits leak target-specific signal and inflate apparent accuracy.

### Verify integrity if in doubt

Decode a graph’s amino-acid and chain channels, re-hash, and confirm the result equals the filename content key; the ESM-2 features re-embed to cosine 1.000. Per-file SHA-256 checksums are in `SHA256SUMS` and a Croissant 1.0 record is in `croissant.json`.

### Intended uses

Benchmarking affinity predictors under leakage-free antigen splits; affinity maturation by measure-then-rank, where a handful of measured variants against a target are used to rank the next round of candidates; and training or pretraining sequence- and structure-based affinity models on a single standardized dataset.

### Not intended for

Zero-shot affinity prediction for antigens absent from the training set; interface-level use of the predicted structures without confidence filtering (interface accuracy is contaminated-optimistic and collapses on novel complexes, Section 4.4); clinical decision-making without experimental validation.

### Known limitations

Affinities aggregate heterogeneous assays (SPR, BLI, ITC, ELISA, high-throughput deep mutational scanning, and curated literature), with inter-source measurement offsets that persist despite unit normalization; the 3D complexes are predictions, not experimental structures; and, as noted in Section 2.2, the public landscape imposes a label-diversity ceiling that limits cross-target prediction.

### Ethics

No human or animal subjects were involved; all records derive from publicly available databases, each retaining its original license (`PROVENANCE.md`).

## 6 Code and Data Availability

### Availability

The dataset (metadata, predicted structures, embeddings, graphs, and documentation) and all construction and evaluation code are deposited in Zenodo (Data Citation 1) with a DOI assigned on deposit, and mirrored in the project repository. The curated compilation and all derived artifacts are released under a Creative Commons Attribution 4.0 (CC BY 4.0) license; individual source datasets retain their original licenses, documented per source in `PROVENANCE.md`. Records from two sources whose licenses do not permit redistribution (AgAb/ASD, non-commercial; Ab-CoV, no explicit reuse license; 327 records) are provided as source pointers to be obtained from the original providers under their terms.

### Reproducibility

The full curation pipeline (aggregation, decontamination, de-placeholding, rebalancing, deduplication, structural enrichment, and content-key regeneration), the feature- and graph-generation scripts, and the evaluation code (antigen-grouped cross-validation) are released as deterministic, parameterized scripts, so the release can be reproduced end-to-end or adapted. Structure prediction uses Boltz-2 [20]; sequence numbering uses ANARCI [19]; representations use ESM-2 [21]; graph construction uses PyTorch Geometric [25] with GVP features [5]; antigen clustering uses MMseqs2 [22].

### Maintenance

GRABS is maintained as versioned, content-addressed releases, each accompanied by a changelog and per-file SHA-256 checksums.

## Author Contributions

L.M. designed and built the dataset, implemented the curation pipeline and validation experiments, and wrote the manuscript. J.M. and P.P.Z. supervised the work and revised the manuscript.

## Data Citations

1. Martins, L., Zenatti, P. P. & Meidanis, J. GRABS: a content-addressed antibody-antigen binding-affinity dataset with predicted structures and a leak-free benchmark (v1.1). *Zenodo* https://doi.org/10.5281/zenodo.24119328 (2026).

